# Comparison of gene set scoring methods for reproducible evaluation of multiple tuberculosis gene signatures

**DOI:** 10.1101/2023.01.19.520627

**Authors:** Xutao Wang, Arthur VanValkenberg, Aubrey R. Odom-Mabey, Jerrold J. Ellner, Natasha S. Hochberg, Padmini Salgame, Prasad Patil, W. Evan Johnson

## Abstract

**Rationale:** Many blood-based transcriptional gene signatures for tuberculosis (TB) have been developed with potential use to diagnose disease, predict risk of progression from infection to disease, and monitor TB treatment outcomes. However, an unresolved issue is whether gene set enrichment analysis (GSEA) of the signature transcripts alone is sufficient for prediction and differentiation, or whether it is necessary to use the original statistical model created when the signature was derived. Intra-method comparison is complicated by the unavailability of original training data, missing details about the original trained model, and inadequate publicly-available software tools or source code implementing models. To facilitate these signatures’ replicability and appropriate utilization in TB research, comprehensive comparisons between gene set scoring methods with cross-data validation of original model implementations are needed.

**Objectives:** We compared the performance of 19 TB gene signatures across 24 transcriptomic datasets using both re-rebuilt original models and gene set scoring methods to evaluate whether gene set scoring is a reasonable proxy to the performance of the original trained model. We have provided an open-access software implementation of the original models for all 19 signatures for future use.

**Methods:** We considered existing gene set scoring and machine learning methods, including ssGSEA, GSVA, PLAGE, Singscore, and Zscore, as alternative approaches to profile gene signature performance. The sample-size-weighted mean area under the curve (AUC) value was computed to measure each signature’s performance across datasets. Correlation analysis and Wilcoxon paired tests were used to analyze the performance of enrichment methods with the original models.

**Measurement and Main Results:** For many signatures, the predictions from gene set scoring methods were highly correlated and statistically equivalent to the results given by the original diagnostic models. PLAGE outperformed all other gene scoring methods. In some cases, PLAGE outperformed the original models when considering signatures’ weighted mean AUC values and the AUC results within individual studies.

**Conclusion:** Gene set enrichment scoring of existing blood-based biomarker gene sets can distinguish patients with active TB disease from latent TB infection and other clinical conditions with equivalent or improved accuracy compared to the original methods and models. These data justify using gene set scoring methods of published TB gene signatures for predicting TB risk and treatment outcomes, especially when original models are difficult to apply or implement.

## INTRODUCTION

Prior to the COVID-19 pandemic, tuberculosis (TB) was the leading infectious cause of death worldwide (1, 2). Approximately 10 million people developed TB, and 1.4 million died from the disease (1). Current diagnostic tests for TB disease include sputum acid-fast bacilli (AFB) smear microscopy, rapid molecular tests, and culture-based technologies (1). With the advent and widespread availability of nucleic acid amplification tests (Xpert MTB/RIF), most cases of pulmonary TB can be diagnosed quickly and accurately. Paucibacillary TB diagnosis (smear-negative pulmonary, extrapulmonary, and pediatric TB) (3–6) and predicting treatment success/failure remain difficult challenges. Furthermore, there are gaps in identifying individuals with quiescent or percolating disease, and utilizing non-sputum-based technologies would facilitate diagnosis for individuals unable to produce sputum (e.g., children). There is therefore an urgent need for additional technologies that ensure high-quality, timely, effective testing for people living with TB (1, 7).

Over the last ten years, multiple blood-based biomarkers using gene expression profiles have been developed for TB. These signatures can distinguish active TB disease from latent TB infection (LTBI) (8, 9), distinguish TB from other diseases (OD) (10–12), predict progression from LTBI to active TB (13, 14), and assess treatment response (15). These or other signatures may potentially meet target product profiles (TPP) proposed by the World Health Organization (WHO) for point-of-care (POC) testing, since some studies have shown that perturbation in the gene expression profile occurs prior to sputum conversion (13, 14, 16, 17). Blood-based signatures could also identify individuals for additional diagnostics or LTBI treatment. However, more research must be done to establish the efficacy and reproducibility of using blood-based signatures in the field, as shown by the CORTIS trial (18).

The common approach to building a gene signature is to identify a *gene set*, a subset of the thousands of genes whose expression is measured for a patient cohort, using a combination of domain knowledge and data-driven feature selection methods. The gene set is then used to train a gene signature using a *prediction algorithm* such as regression or some other machine learning approaches for deciding how to turn gene expression measurements into an outcome prediction. Typically, biomarkers are *trained* using one subset of data and then finalized or *fixed* before being applied for prediction in new samples and cohorts. However, in the case of existing TB signatures, the *replicability* of these biomarkers is inadequate, meaning that many of the original publications did not give enough detail to replicate the published models. In addition, nearly all of these publications failed to provide usable code to rerun the published model on the study data—let alone apply the models to new datasets. Some of these original models further lacked *reproducibility* of the performance and accuracy of TB gene signatures—meaning that the signatures were *overfit* and thus experienced significant reductions in performance in later observations compared to the training set (10, 19, 20). Many factors likely contribute to this phenomenon, including overfitting to encodes in the training data, incomplete validation of the biomarker, differences in study population demographics or comorbidities, and possible inconsistencies due to differences in the platforms used to measure gene expression (37, 46).

Several research teams have attempted to address issues of reproducibility in TB gene expression biomarker research, either by rebuilding the original classification models (7, 19) or by using methods such as gene set enrichment analysis (GSEA) in place of the original models (21). For example, Warsinske et al. (19) reconstructed the original models for 16 TB gene signatures and compared gene signature accuracy in the contexts of TB diagnostics and progression to disease. For some of their biomarkers, the discovery dataset(s) used to train the gene signatures were unavailable, or the proposed diagnostic models comprised the gene signatures were difficult to reproduce. Other challenges included limited information about data preprocessing and hyperparameter values used to build the model (22–24). Despite clear documentation of their reconstruction of original models, Warsinske et al. (19) did not publish software code for their work from scratch to use the biomarkers from their comparisons.

As a result of this deficit in replicability, we have endeavored to fill this gap. Our team has recently developed and released the open-source TBSignatureProfiler software toolkit, which provides a compilation of TB gene sets used from published biomarkers and provides methods (21) to evaluate the performance of existing TB gene sets on new datasets via the application of various scoring algorithms (25) (e.g., GSVA, ssGSEA, PLAGE). However, while alternative methods, such as gene set scoring are simpler to use compared to the original models, these methods have not been established as reasonable approximations or alternatives to original model performance.

To address the issues of reproducibility in reconstructing the discovery set, our study uniformly evaluated the performance of 19 TB gene signatures across 24 datasets using both original models and gene set scoring methods through the TBSignatureProfiler platform. We systematically compared our results to previous research (19) to the signature’s ability to distinguish patients with active TB from other clinical conditions. Our results illustrated that gene set scoring methods are effective computational tools for profiling the performance of TB gene signatures and produced similar accuracy compared to the original model, demonstrating alternative, accessible ways to profile existing TB gene signatures. We also sought to facilitate the replicability of TB gene signatures’ application and comparative analysis. We therefore curated the TB transcriptomic datasets used in this study and included the corresponding discovery model for each gene signature in the TBSignatureProfiler R package, enabling the reproducibility of all results.

## METHODS

### Gene signatures and original diagnostic models used for comparison

Nineteen existing TB gene signatures were selected for this study primarily based on the results of Warsinske *et al*. (Table 1) (10, 12–14, 22–24, 26–32) to make a fair comparison for the performance of these signatures. Sixteen gene signatures were trained to predict active TB versus other clinical conditions. Three additional gene signatures that predict progression from LTBI to active TB disease, Zak_RISK_16 (14), Suliman_RISK_4 (13), and Leong_RISK_29 (32), were also included. Twenty-four datasets (9–12, 16, 21, 27–29, 31, 33–48) that include subjects with active TB and other clinical conditions have been selected for evaluation. These datasets were selected in alignment with Warsinske *et al*. to systematically compare different gene scoring methods and the original models (19). Twenty of these datasets were from microarray studies; the other four consisted of RNA-sequencing data. A detailed description of the transcriptomic studies included in this study can be found in the Supplementary Material.

**Table 1.** Summary of TB gene signatures compared in this study and their original training model.

| Signature Name | Gene Signature Number | Comparison | Datasets | Original Model Description |
| --- | --- | --- | --- | --- |
| Sweeney_OD_3 | 3 | Active tuberculosis vs. (LTBI & HCs & OD) | GSE19491 & GSE42834 & GSE37250 | Difference of geometric means between up and downregulated genes |
| Jacobsen_3 | 3 | Active tuberculosis vs. LTBI | GSE19491 | Linear Discriminant Analysis |
| LauxdaCosta_OD_3 | 3 | Active tuberculosis vs. OD | GSE42834 | Random Forest |
| Maertzdorf_4 | 4 | Active tuberculosis vs. HCs | GSE74092 | Random Forest |
| Sambarey_HIV_10 | 10 | Active tuberculosis vs. OD | GSE37250 | Linear Discriminant Analysis |
| Verhagen_10 | 10 | Active tuberculosis vs. (LTBI & HCs) | GSE41055 | Random Forest |
| Maertzdorf_15 | 15 | Active tuberculosis vs. HCs | GSE74092 | Random Forest |
| Leong_24 | 24 | Active tuberculosis vs. LTBI | GSE10175 | Ridge Logistic Regression |
| Kaforou_27 | 27 | Active tuberculosis vs. OD | GSE19491 | Difference of arithmetic means between up and downregulated genes |
| Anderson_42 | 42 | Active tuberculosis vs. LTBI | GSE39940 | Difference of sums between up and downregulated genes |
| Kaforou_OD_44 | 44 | Active tuberculosis vs. OD | GSE19491 | Difference of arithmetic means between up and downregulated genes |
| Anderson_OD_51 | 51 | Active tuberculosis vs. OD | GSE39940 | Difference of sums between up and downregulated genes |
| Kaforou_OD_53 | 53 | Active tuberculosis vs. OD | GSE19491 | Difference of arithmetic means between up and downregulated genes |
| Berry_OD_86 | 86 | Active tuberculosis vs. OD | GSE19491 | K-nearest neighbors algorithm |
| Bloom_OD_144 | 144 | Active tuberculosis vs. (HCs & OD) | GSE42834 | Support Vector Machines |
| Berry_393 | 393 | Active tuberculosis vs. (LTBI & HCs) | GSE19491 | K-nearest neighbors algorithm |
| Suliman_RISK_4 | 4 | Incipient tuberculosis vs. HCs | GSE94438 | Support Vector Machines (linear kernel, using paired ratio) |
| Zak_RISK_16 | 16 | Incipient tuberculosis vs. HCs | GSE79362 | Support Vector Machines (linear kernel) |
| Leong_RISK_29 | 29 | Incipient tuberculosis vs. HCs | GSE79362 | Lasso Logistic Regression |

The reconstruction of each signature’s original model followed the guidelines given by Warsinske et al. (20), who reconstructed these models based on information from each of the signature’s original publications. Two training methods were used for TB gene signature discovery: model-based and score-based approaches. The model-based approaches use machine learning algorithms such as support vector machines (33) or random forests (49) to reconstruct the biomarker’s original model. The score-based approaches primarily rely on the difference of sums or arithmetic means between the upregulated genes and the downregulated genes within the gene set and therefore do not require model retraining. Six of the 19 identified gene signatures used score-based methods to distinguish active TB patients from other clinical conditions (Table 1). The reconstructed models were validated against the performance reported in the original publication using the AUC metric. To evaluate a biomarker’s ability to identify patients with active TB, the associated model was applied across the 24 curated transcriptomic studies.

### Gene set scoring methods

We used single sample GSEA (ssGSEA) (50), Gene Set Variation Analysis (GSVA) (35), Pathway level analysis of gene expression (PLAGE) (51), Zscore (52), and Singscore (unidirectional and bidirectional versions) (50, 53) to evaluate the accuracy of TB gene signatures across 24 studies. Please see the Discussion section for a more detailed comparison of these five gene set scoring methods. The evaluation of distinct gene signatures using different gene set scoring methods and its original model was performed by the runTBSigProfiler function from the TBSignatureProfiler R package (38).

### Gene signature splitting strategy

Some gene set scoring methods, including GSVA, ssGSEA, and Singscore, rank a signature’s genes against other genes in the dataset. Thus, gene sets with both upregulated and downregulated genes do not usually score well with these methods— they effectively counteract each other in the score computation. Most of the 19 signatures contain genes expressed strongly in both directions; thus, we proposed a ‘signature splitting’ strategy for gene signatures with equal to or more than ten genes to overcome this limitation. Gene signatures with less than ten genes were excluded here as their upregulated or downregulated gene subset was either empty or only contained one gene. In such cases, signature splitting would have a decreased effect on these gene sets compared to a large set.

For the ssGSEA and GSVA methods, we evaluated each signature’s performance using only the upregulated or the downregulated subsets of genes within the gene sets. For gene signatures that did not require reconstruction of the training model, the upregulated and downregulated genes were usually provided by the original publication. If unavailable in the original publication, the upregulated or the downregulated genes were identified by performing differential expression analysis on the datasets from which the associated biomarkers were derived using the R package DESeq2 (54) for RNA-seq data and limma (55) for microarray data. For the bidirectional implementation of Singscore, we used the simpleScore function from the singscore R package (53) to algorithmically combine the disease score for the upregulated and downregulated genes.

### Imputation of missing genes and removal of batch effects

Occasionally, various genes within gene sets were missing across transcriptomic studies, mainly due to the inconsistent gene coverage across platforms or different versions of the gene symbol naming systems across different sequencing platforms. This might affect a biomarker’s predictive accuracy in some datasets and across biomarker methods. Two steps were taken to mitigate this issue. First, we updated gene symbols for gene signatures and candidate datasets using the HGNChelper R package (56) to reduce gene misidentification during the training and validation process. More details on the matching of probe ID to gene symbol and missing gene information within each dataset can be found in the Supplementary Material. Second, a k-nearest neighbors (KNN) algorithm was applied to impute the missing gene expression values based on the signature’s discovery dataset(s) (57). This procedure may elicit potential bias problems when large portions of genes within a signature are missing in some datasets. To make a fair comparison, we applied this imputation procedure to all biomarkers that used model-based diagnostic approaches across independent datasets. Functionality *reference ComBat* (58, 59) was used to make generalized comparisons of signatures across the independent studies. *reference ComBat* removes batch effects between the training datasets and the testing datasets without changing the reference dataset’s gene expression values. With this functionality, the biomarker’s original model does not have to be retrained whenever the batch correction is performed or when the model is applied to a new dataset, increasing efficiency when evaluating biomarkers with model-based approaches.

### Statistical Evaluation of Model Performance

The AUC value corresponding to sample scores against disease subtypes for each TB gene signature was calculated for each dataset. The sample-size-weighted mean AUC (weighted AUCs) was used to assess the overall performance of each gene set across independent studies while excluding the discovery dataset(s) used to train the corresponding signature (44). AUC results were calculated with the R package ROCit (60) compute estimates. Each signature’s 95% confidence interval for the weighted mean AUC value was calculated using the bootstrap method with 10,000 bootstrap resamples. The Wilcoxon paired test was applied to test whether AUCs given by different methods were significantly different from each other.

### Evaluating biomarker performance based on original models

The original models were inputted into the TBSignatureProfiler (45) platform, which provides a computational profiling platform that allows researchers to evaluate multiple TB gene signatures. In addition to the original models, users can access other scoring methods and gene sets in the TBSignatureProfiler. The TBSignatureProfiler was used to conduct all the analyses for this work, and the package can be downloaded from GitHub (https://github.com/compbiomed/TBSignatureProfiler) or Bioconductor (http://www.bioconductor.org/packages/release/bioc/html/TBSignatureProfiler.html). Code for the analyses from this paper can be found at: https://github.com/xutao-wang/Comparison-of-existing-tuberculosis-gene-signatures.

### Comparing the performance of gene set scoring methods with original models

We noticed that there are several studies (i.e. GSE81746, GSE107331, GSE29536 etc.) with a small sample sizes (n < 40) which only consisted of healthy controls (HCs) and subjects with active TB. For these studies, most was generalizable to gene signatures that could distinguish patients with active TB phenotype with high predictive accuracy; however, such performance could not be generalized toward large populations or to cohorts with heterogeneous disease types including LTBI. To produce a fair comparison of gene signatures’ performance using gene set scoring methods and their original models, for each profiling method (including both gene set scoring methods and original models) we selected datasets with a sample size greater than 40 and AUC estimates greater than 0.80 across all TB biomarkers simultaneously excluding the corresponding training study for each biomarker.

Several metrics were used to compare the performance of gene signatures as assessed by different gene set scoring methods and their original models. For each TB gene signature, the Spearman’s rank correlation (*ρ*) was computed to measure the strength of association of the prediction scores from a signature’s original model and the different gene set scoring methods. We then summarized correlation results, computing the weighted Spearman’s rank correlation (*ρ* _*w*_), based on equation 1.

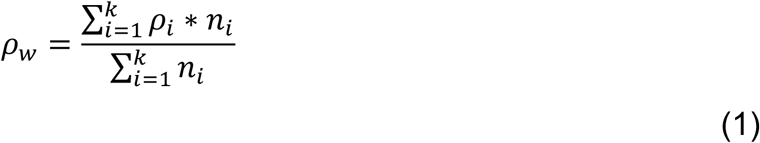

Moreover, the absolute AUC difference (|∆*AUC*|) between the original model and various gene set scoring methods were computed as shown in equation 2.1 for each selected dataset, with the weighted absolute AUC difference |∆*AUC*|_*w*_ representing the overall distribution pattern across all selected studies (equation 2.2). Additionally, the Szymkiewicz–Simpson coefficient (61) or overlap coefficient (*oc*) as shown by equation (3), was applied to evaluate the similarity of studies based on the results given by each of the gene set scoring method and original model for all gene signatures.

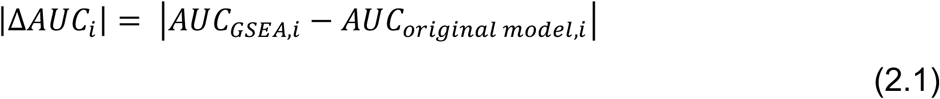

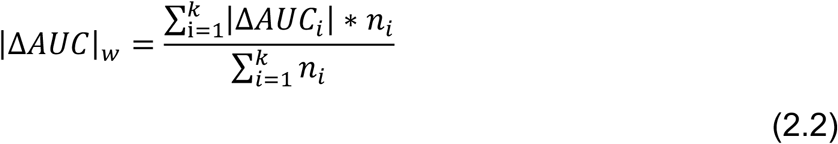

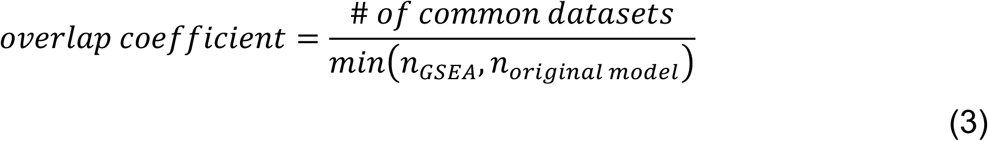

## RESULTS

### Evaluation of discovery studies using training models from original publications

The list of signatures and information on their respective training data are presented in Table 1, along with AUC values from their originating publication, Warsinske *et al*., and our results. The AUC values for our reconstructed models were nearly identical to the results of the original publications, suggesting that the training models were accurately reconstructed. Several gene signatures including Sweeney_OD_3, Maertzdorf_15, Leong_24, Kaforou_27, Anderson_42, and Berry_393, when evaluated by ssGSEA, had estimated AUC values above 0.9 (Table S1). These results suggest that ssGSEA is a comparable signature profiling method, producing accurate results for some TB signatures in differentiating the TB disease states (active TB vs. LTBI, HCs, or OD) even if the evaluation studies were the signature’s discovery datasets.

### Performance of original models and gene set scoring methods for active TB versus non-active TB

Figure 1 shows the AUC distribution of 24 independent TB studies across 19 TB gene signatures based on the results from their original models. 13 of 16 signatures had AUCs greater than 0.9 from their discovery dataset(s) (Figure 1). Each TB signature’s original model was applied across all the datasets, and their performance varied greatly on a study-by-study basis (Figure 1). Notably, the Kaforou_OD_53, Kaforou_27, Maertzdorf_15, and Sweeney_OD_3 signatures had consistently high AUC values across different studies, with weighted AUCs of 0.83 for Kaforou_OD_53, 0.81 for Kaforou_27, 0.82 for Maertzdorf_15, and 0.81 for Sweeney_OD_3 (Table 2). In contrast, the performance of some signatures, such as Verhagen_10, was mixed across studies (Figure 1). Verhagen_10 had a weighted mean AUC of 0.61 (Table 2), performing well in some datasets (>0.9 AUC in GSE81746, GSE41055, and GSE29536) but with poor performance in most of the remaining studies (<0.65 AUC in 16 out of 24 studies; Figure 1). Noted that Zak_RISK_16, Suliman_RISK_4, and Leong_PREDICT_29 also performed poorly in these comparisons, but these are signatures of disease progression (Figure 1).

**Table 2.** Weighted mean AUC and 95% CI for 19 gene signatures using original model and gene set scoring methods (ssGSEA, GSVA, PLAGE, Zscore, and Singscore) across 24 studies. Color-coded for AUC values that were 0.05 unit greater (red) than in Warsinske et al and were 0.05 less (blue) than Warsinske et al. (*: p <= 0.01, **: p <= 0.001, ***: p <= 0.0001 derived from Wilcoxon signed-rank test between the original model and corresponding gene set scoring methods; †: the original model results underperformed the Warsinske et al. with more than 0.05 unit; ‡: the original model results outperformed the Warsiske et al. with more than 0.05 unit).

| Signature | Warsinske et al. | TBSignatureProfiler<br>(Original Model) | TBSignatureProfiler<br>(ssGSEA) | TBSignatureProfiler<br>(GSVA) | TBSignatureProfiler<br>(PLAGE) | TBSignatureProfiler<br>(Zscore) | TBSignatureProfiler<br>(Singscore) |
| --- | --- | --- | --- | --- | --- | --- | --- |
| LauxdaCosta_OD_3 | 0.76 (0.45–1.00) | 0.80 (0.76-0.83) | 0.82 (0.76-0.86) | 0.79 (0.74-0.83) | 0.83* (0.79-0.86) | 0.78 (0.74-0.80) | 0.82 (0.75-0.86) |
| Jacobsen_3 | 0.83 (0.69–0.98) | 0.76† (0.71-0.80) | 0.72 (0.68-0.76) | 0.69 (0.64-0.73) | 0.78 (0.75-0.81) | 0.69 (0.65-0.73) | 0.71 (0.67-0.75) |
| Sweeney_OD_3 | 0.85 (0.72–0.99) | 0.81 (0.76-0.85) | 0.82 (0.77-0.86) | 0.75 (0.69-0.80) | 0.82 (0.78-0.85) | 0.77 (0.71-0.82) | 0.82 (0.77-0.86) |
| Maertzdorf_4 | 0.79 (0.64–0.95) | 0.80 (0.77-0.83) | 0.70* (0.65-0.75) | 0.73** (0.69-0.78) | 0.81 (0.78-0.85) | 0.72** (0.68-0.76) | 0.66* (0.62-0.72) |
| Verhagen_10 | 0.54 (0.41–0.68) | 0.61‡ (0.57-0.66) | 0.57 (0.55-0.60) | 0.59 (0.56-0.64) | 0.65 (0.59-0.71) | 0.62 (0.57-0.68) | 0.58 (0.55-0.61) |
| Sambarey_HIV_10 | 0.82 (0.57–1.00) | 0.83 (0.76-0.87) | 0.80 (0.76-0.84) | 0.80 (0.76-0.84) | 0.76 (0.70-0.83) | 0.75 (0.70-0.81) | 0.79 (0.75-0.83) |
| Maertzdorf_15 | 0.79 (0.66–0.92) | 0.82 (0.79-0.85) | 0.83 (0.79-0.87) | 0.82 (0.78-0.86) | 0.83 (0.80-0.86) | 0.79 (0.75-0.82) | 0.79 (0.75-0.84) |
| Leong_24 | 0.75 (0.54–0.95) | 0.72 (0.67-0.78) | 0.73 (0.70-0.77) | 0.61 (0.58-0.65) | 0.73 (0.67-0.79) | 0.61 (0.58-0.65) | 0.73 (0.69-0.78) |
| Kaforou_27 | 0.83 (0.64–1.00) | 0.81 (0.77-0.85) | 0.82 (0.78-0.85) | 0.79 (0.76-0.82) | 0.83 (0.79-0.87) | 0.79 (0.76-0.82) | 0.78 (0.73-0.82) |
| Anderson_42 | 0.82 (0.66–0.97) | 0.78 (0.73-0.83) | 0.61** (0.58-0.66) | 0.60** (0.58-0.64) | 0.76 (0.70-0.82) | 0.65** (0.59-0.73) | 0.57** (0.53-0.61) |
| Kaforou_OD_44 | 0.78 (0.56–1.00) | 0.76 (0.69-0.81) | 0.67 (0.63-0.71) | 0.72 (0.68-0.74) | 0.80 (0.72-0.86) | 0.70 (0.67-0.74) | 0.70 (0.67-0.74) |
| Anderson_OD_51 | 0.58 (0.33–0.82) | 0.71‡ (0.64-0.78) | 0.75 (0.71-0.80) | 0.81 (0.77-0.84) | 0.79 (0.72-0.85) | 0.75 (0.66-0.82) | 0.77 (0.73-0.80) |
| Kaforou_OD_53 | 0.84 (0.70–0.99) | 0.83 (0.78-0.87) | 0.70** (0.66-0.75) | 0.77** (0.74-0.80) | 0.84 (0.80-0.87) | 0.77** (0.73-0.80) | 0.77** (0.73-0.81) |
| Berry_OD_86 | 0.69 (0.36–1.00) | 0.69 (0.66-0.72) | 0.71 (0.68-0.76) | 0.74 (0.70-0.78) | 0.75* (0.72-0.80) | 0.73 (0.69-0.78) | 0.73 (0.68-0.78) |
| Bloom_OD_144 | 0.74 (0.52–0.96) | 0.70 (0.66-0.74) | 0.77 (0.72-0.81) | 0.76 (0.71-0.81) | 0.71 (0.66-0.78) | 0.70 (0.63-0.77) | 0.76 (0.71-0.81) |
| Berry_393 | 0.71 (0.43–0.99) | 0.70 (0.66-0.74) | 0.78 (0.74-0.82) | 0.77 (0.73-0.81) | 0.79* (0.75-0.84) | 0.77 (0.73-0.81) | 0.79 (0.74-0.84) |
| Suliman_RISK_4 | NA | 0.62 (0.57-0.69) | 0.62 (0.58-0.68) | 0.55 (0.53-0.58) | 0.74** (0.70-0.79) | 0.60 (0.55-0.66) | 0.61 (0.56-0.66) |
| Zak_RISK_16 | NA | 0.62 (0.56-0.70) | 0.85*** (0.81-0.88) | 0.84*** (0.80-0.88) | 0.83*** (0.80-0.86) | 0.83*** (0.79-0.86) | 0.84*** (0.80-0.88) |
| Leong_RISK_29 | NA | 0.68 (0.65-0.73) | 0.58* (0.56-0.61) | 0.59* (0.56-0.61) | 0.75 (0.70-0.80) | 0.60 (0.56-0.64) | 0.61* (0.57-0.65) |

**Figure 1.**
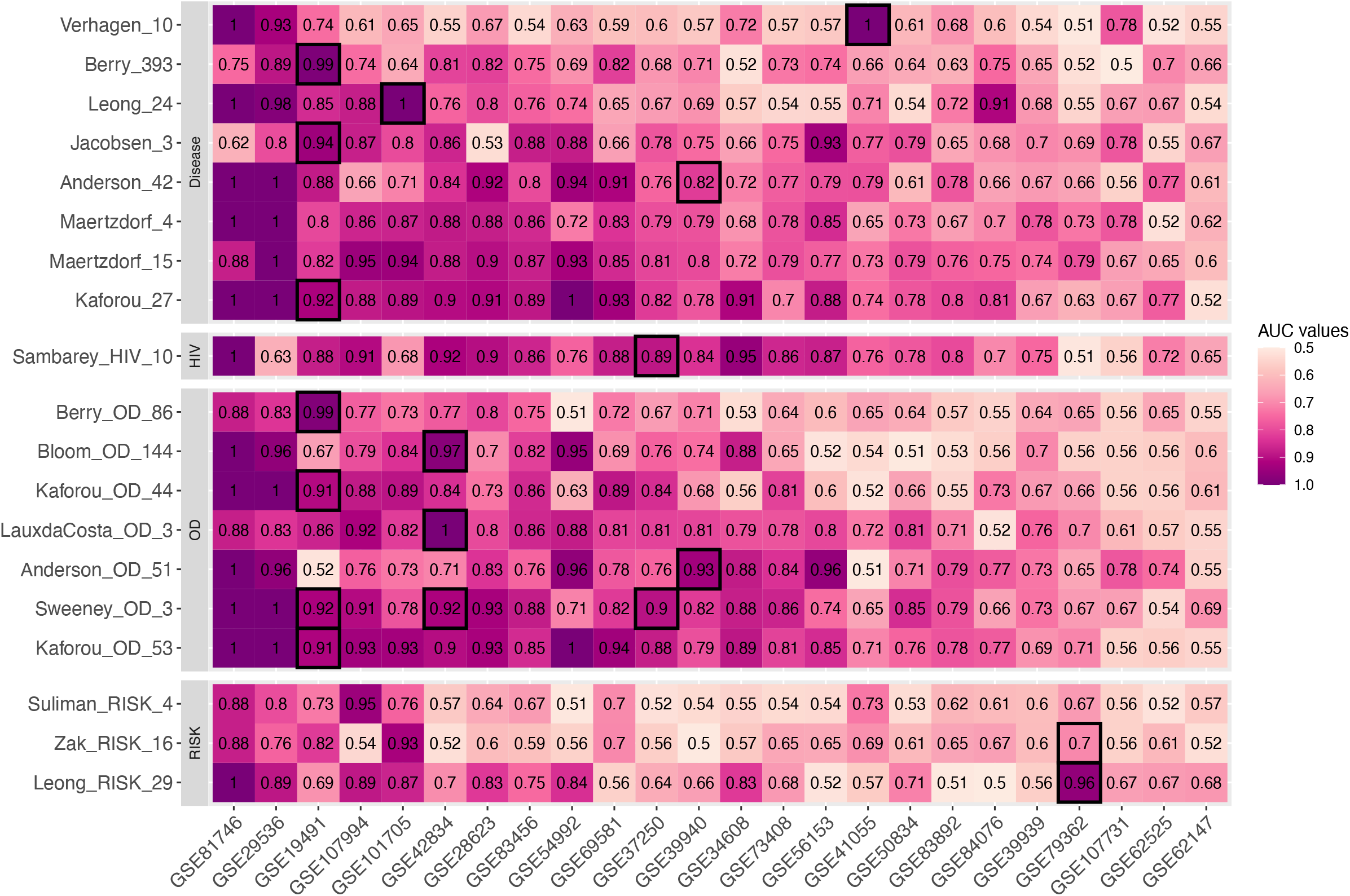
Heatmap of AUC distribution for each signature across 24 studies using original model. Grids with black borders were the discovery sets for each TB gene signature. Each row represents one signature. Signatures were clustered into different categories with respect to the TB subtypes that they aim to identify. The column of the heatmap representing the studies that were used in this paper. The datasets were rearranged in decreasing order based on their mean AUC values across all TB gene signatures.

Table 2 shows the weighted AUCs for each gene signature as computed using their corresponding original model and five different gene set scoring methods. In this table, AUC values in red denote results from our reimplemented models and gene set scoring methods that outperformed the original model with differences greater than 0.05 units. Values in blue represent cases where AUC values from the original model outperformed other methods the methods by more than 0.05 units. The 0.05 threshold was chosen mainly for visualization purposes, which suggested that there was a noticeable difference in the weighted AUCs given by two different methods.

16 of 19 gene signatures had reimplemented weighted AUCs within 0.05 units compared to Warsinske *et al*. (Table 2). However, our reimplemented models for Verhagen_10 and Anderson_OD_51 outperformed the results from Warsinske *et al*., and our original model for Jacobsen_3 underperformed compared to Warsinske *et al*. (Table 2). The weighted AUCs for Anderson_OD_51 computed by gene scoring methods surpassed that of its original model but not the results given by ssGSEA, although none of the results were statistically significant after adjusting for multiple testing (p-value > 0.01; Table 2). For Berry_393, the weighted AUCs computed from its original model underperformed all the five gene scoring methods; specifically, the PLAGE AUC was 0.79 (95% CI: 0.75–0.84), which was significantly higher (p-value <= 0.01) than that of the original model which had an AUC of 0.70 (0.66–0.74). Results from Zak_RISK_16 given by its original model also underperformed five gene scoring methods, and all the superior weighted AUCs were significant (p-value <= 1e-04) (Table 2). Additionally, the performance of each gene signature across individual datasets using five gene set scoring methods was presented in Figure S1A-E.

Figure 2A shows the weighted mean AUC difference between the original model and the corresponding five gene scoring methods across all gene sets. The outperformance of the weighted AUCs given by PLAGE was consistent in most of the gene signatures asides from Sambarey_HIV_10 and Anderson_42. The accompanying Wilcoxon signed-rank test results indicated that the results from PLAGE were significantly different from the results of the original models (p-value <= 0.01), whereas the results from the remaining four gene scoring methods were statistically equivalent when compared to the weighted AUCs given by the original model (p-value > 0.05; Figure 2A).

**Figure 2.**
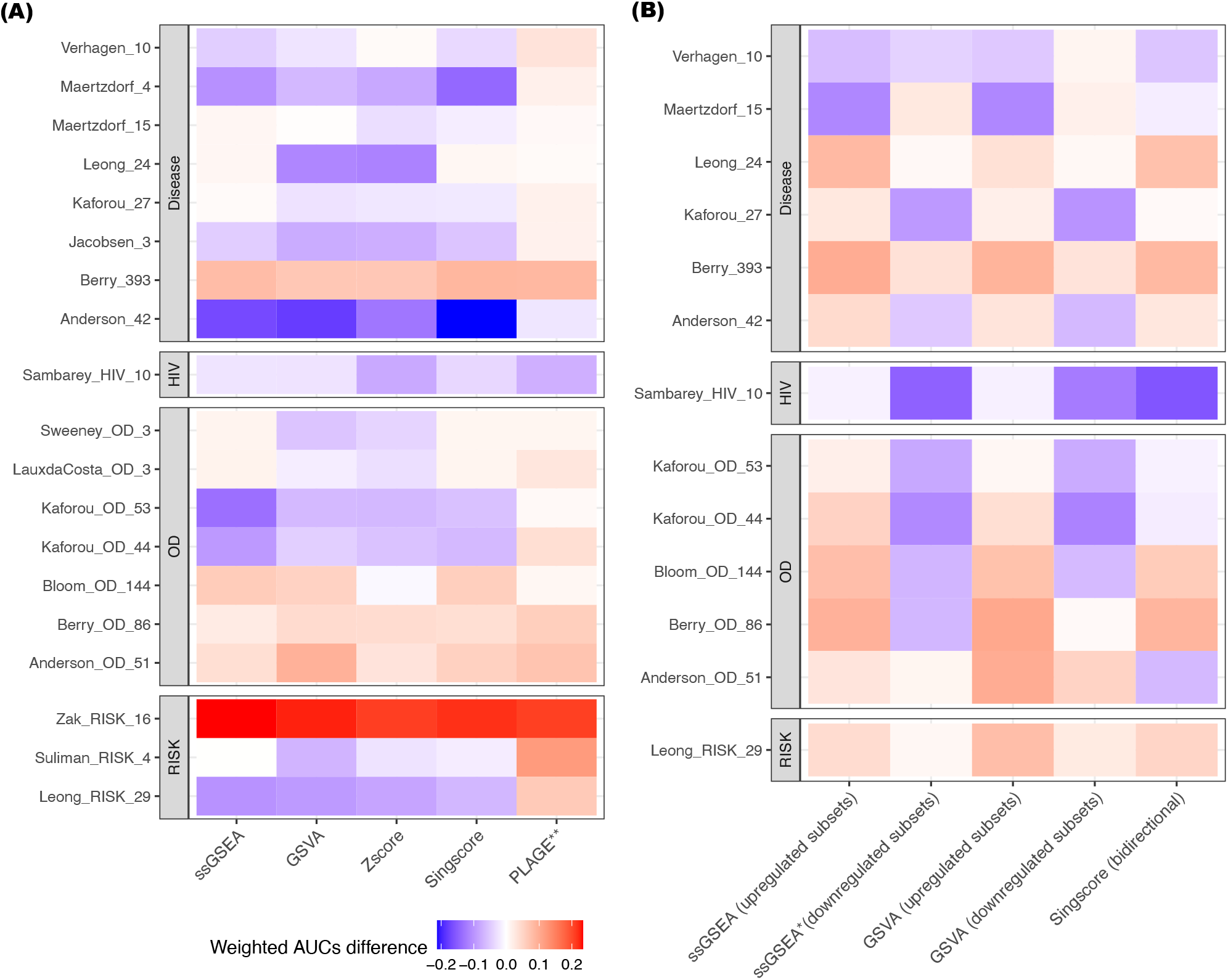
Differences of the weighted mean AUC values between the original model versus gene set scoring methods. Each grid showed the difference of weighted mean AUC value between the use of the corresponding gene set scoring methods (ssGSEA, GSVA, PLAGE, and Singscore) and the use of original models. The results from the original model was used as the baseline. For all 19 TB gene signatures, the weighted mean AUC results computed by ssGSEA, Singscore, GSVA, and Zscore were statistically equivalent from the results given by the original model **(A)**. The weighted mean AUC results computed by ssGSEA and GSVA for the upregulated subsets of the gene signatures, Singscore (bidirectional scoring), and GSVA for the downregulated subsets of the gene signatures were statistically equivalent from the results given by the original model **(B)**. Red: the weighted mean AUC for gene scoring method outperformed the original model. Blue: the weighted mean AUC for gene scoring method underperformed the original model. *: p-value < 0.05 derived from the Wilcoxon signed-rank test **: p-value < 0.01 derived from Wilcoxon signed-rank test

#### Accounting for gene expression direction

Table 3 shows the weighted mean AUCs from ssGSEA and GSVA after splitting the genes into upregulated or downregulated subsets, along with the Singscore bidirectional scoring results. There was significant improvements in the weighted AUCs for the upregulated subsets of Berry_OD_86 based on the ssGSEA (AUC = 0.78; 95% CI: 0.75–0.82) and GSVA (AUC = 0.80; 95% CI: 0.76–0.84) results when compared to its original models (p-value <= 1e-04 for both methods; Table 3). Similar improvement was observed when using ssGSEA and GSVA for the upregulated subsets of Berry_393 (p-value <= 0.01 for both methods). Interestingly, Anderson_OD_51 is the only gene signature for which the weighted AUCs from GSVA (upregulated subsets: AUC = 0.81, 95% CI: 0.78-0.84; p-value = 0.03; downregulated subsets AUC = 0.77, 95% CI: 0.71-0.82; p-value = 0.04) outperformed its original model (Table 3). Lastly, the performance of Berry_OD_86 and Berry_393 given by Singscore bidirectional scoring also outperformed their original models (p-value <= 1e-3 for both gene sets; Table 3). Most gene signatures had similar results when using the unidirectional and bidirectional versions of Singscore, although Sambarey_HIV_10 and Anderson_OD_51 showed significantly lower weighted AUCs using Singscore bidirectional scoring (p-value < 0.01 for both gene sets; Tables 2 and 3).

**Table 3.**
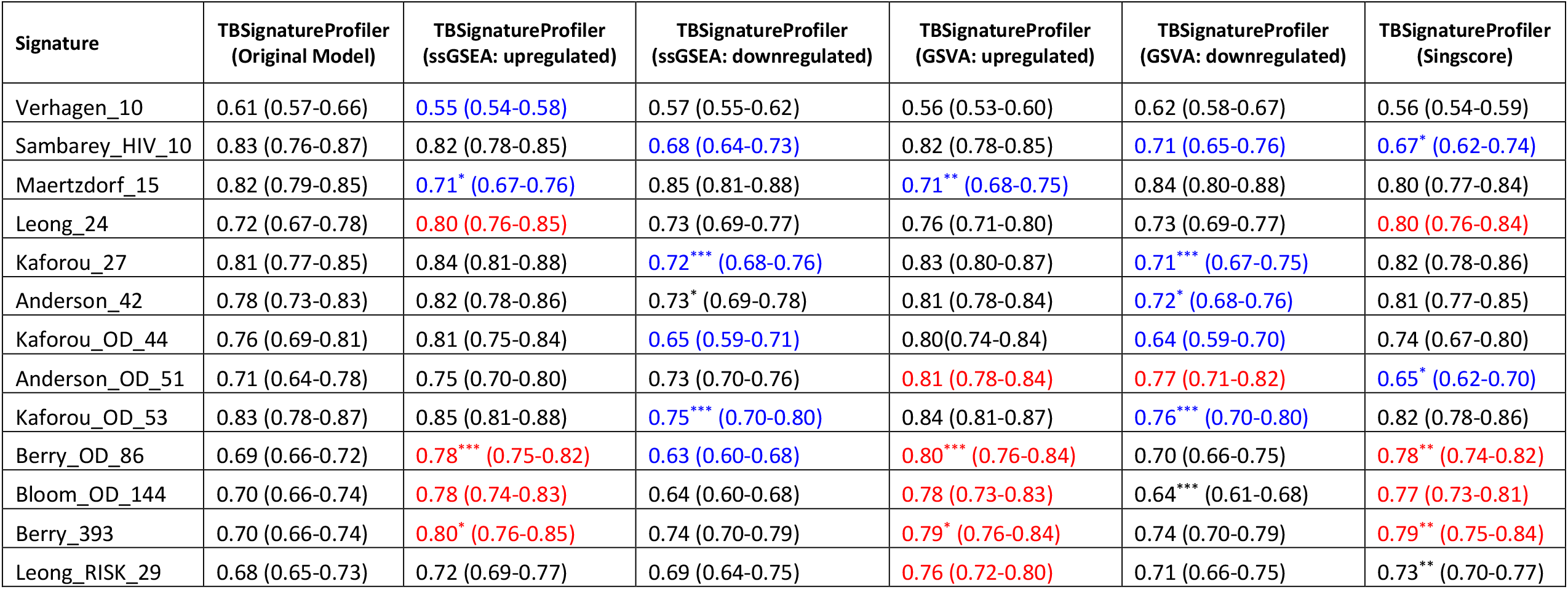
Weighted mean AUC and 95% CI results for upregulated and downregulated subsets of gene signature using the ssGSEA, GSVA, and Singscore (bidirectional version). Color-coded for AUC values that were 0.05 greater (red) than the original model and were 0.05 less (blue) than the original model. (*: p <= 0.01, **: p <= 0.001, ***: p <= 0.0001 derived from Wilcoxon signed-rank test).

Figure 2B illustrates the weighted mean AUC differences between each gene scoring method and original models across gene signatures. Again, the Wilcoxon signed-rank test suggests that the weighted mean AUC results from all methods were statistically equivalent (p-value > 0.05) to the results given by the original model, except for the results computed from the signatures’ downregulated subset using ssGSEA (p-value = 0.036; Figure 2B). Additionally, Figure S2A-C shows the AUC distribution for the up/down-regulated subset of each qualified gene signature across all datasets using the gene set scoring methods ssGSEA, GSVA, and Singscore bidirectional scoring.

Finally, ridge plots displayed the AUC distribution across 24 studies using original models and all the five gene set scoring methods (Figure S3A-C). The gene signatures were ranked in increasing order based on their median AUC values over the individual training dataset used by the given original model. Signatures including Kaforou_27, Kaforou_OD_53, and Sambarey_HIV_10 had statistically equivalent AUC distributions for results obtained from their respective original models, PLAGE, and ssGSEA with the upregulated subset (Kaforou_27 p-value = 0.54, Kaforou_OD_53 p-value = 0.93, Sambarey_HIV_10 p-value = 0.88; Figure S3A).

### Gene set scoring methods versus original models

Figure S4A-E shows the performance of 19 gene signatures after selection, as based on the criteria described in the methods section. Each point represents a single study; the y-axis gives Spearman’s rank correlation of the predicted scores for the original model and respective gene set scoring method, and the x-axis gives the AUC difference between the results given by two different methods. As expected, Verhagen_10 had poor performance when evaluated using its original model, Zscore, and Singscore.

None of the studies had AUCs greater than 0.8 after selection (Figure S4A-E). GSE101705 was the only study selected, but it showed a distinct prediction pattern given by ssGSEA/GSVA when compared to its original model (0.18 <= |∆*AUC*| <= 0.21; 0.35 <= *ρ*<= 0.57). Similar mediocre performance was observed for Suliman_RISK_4, where GSE107994 was the only study selected by ssGSEA and its original model (|∆*AUC*| = 0.061; *ρ*= 0.7; Figure S4A), and no studies were selected using GSVA and Zscore (Figure S4C-D). In addition, no studies had AUCs greater than 0.8 for Leong_RISK_29 when evaluated using ssGSEA, GSVA, and Zscore (Figure S4A; Figure S4C-D)

#### ssGSEA vs. Original Model

Kaforou_27, Maertzdorf_15, and Sweeney_OD_3 showed high diagnostic accuracy in similar studies (*oc* for Kaforou_27 and Maertzdorf_15: = 1.00, *oc* for Sweeney_OD_3: = 0.86) with small AUC differences (p-value > 0.28 for three gene sets) given by ssGSEA and their original models. Significantly positive Spearman’s rank correlations of the predicted scores were shown for Kaforou_27 (*ρ*_*w*_ = 0.78) and Sweeney_OD_3 (*ρ*_*w*_ = 0.86), and significantly negative correlations were produced for Maertzdorf_15 (*ρ*_*w*_ = −0.81; p-value <= 1e-04 for three signatures; Figure S4A). Although more studies with high AUCs were selected for Anderson_OD_51, Berry_393, Berry_OD_86, Bloom_OD_144, Leong_24, and Zak_RISK_16 using ssGSEA, results from these six signatures showed distinct predicted results when compared to their original models (0.032 <= *ρ*_*w*_ <= 0.60; |∆*AUC*|_*w*_ > 0.050; Figure S4A). Conversely, more datasets were selected for Kaforou_OD_44, Kaforou_OD_53, LauxdaCosta_OD_3, and Maertzdorf_4 based on the results from these signatures’ original models (0.070 <= *ρ*_*w*_ <= 0.72, |∆*AUC*|_*w*_ > 0.050; Figure S4A).

#### PLAGE vs. Original Model

For gene sets Kaforou_27, Kaforou_OD_53, LauxdaCosta_OD_3, Maertzdorf_4, and Maertzdorf_15, PLAGE results were nearly equivalent to the results from their original models (Figure S4B). The study overlap coefficient was 1.00 for all five gene signatures, asides from Kaforou_OD_53, with an AUC value of 0.85 (Figure S4B). Specifically, Kaforou_27, Kaforou_OD_53, and LauxdaCosta_OD_3 had significantly negative Spearman’s rank correlations for the predicted results given by two methods (−0.92 < *ρ*_*w*_ < −0.82; p-value < 1e-04 for all three gene sets), with an absolute weighted AUC difference less than 0.030 for all three gene sets. Two Maertzdorf signatures had strong positive Spearman’s rank correlations of the prediction scores and small AUC differences (p-value for Maertzdorf_4: = 0.06, p-value for Maertzdorf_15: = 0.02), with *ρ*_*w*_ of 0.88 for Maertzdorf_4 and of 0.94 for Maertzdorf_15 (p-value < 1e-05; Figure S4B). Asides from these gene sets, 11 of 14 signatures (excluding Anderson_42, Kaforou_OD_44, and Sambarey_HIV_10) had a higher number of studies with high AUCs selected by PLAGE but showed large absolute AUC differences (|∆*AUC*|_*w*_ > 0.080) and weak to moderate correlations (−0.68 <= *ρ*_*w*_ <= 0.62) of the profiling scores when compared to the results from their original methods (Figure S4B).

#### GSVA vs. Original Model

When comparing the results given by GSVA and signatures’ original models, more studies were selected for Anderson_OD_51, Berry_393, Berry_OD_86, Bloom_OD_144, and Zak_RISK_16 with weighted absolute AUC differences all greater than 0.050 and weak Spearman’s rank correlation (0.11 <= *ρ*_*w*_ <= 0.47) of the predicted scores (Figure S4C). Similar to that of ssGSEA, the resulting scores from Maertzdorf_15 (*ρ*_*w*_ = −0.78, |∆*AUC*|_*w*_ = 0.043) and Maertzdorf_4 (*ρ*_*w*_ = −0.63, |∆*AUC*|_*w*_ = 0.051) were negatively correlated with the results from the signatures’ original model, with a study overlap coefficient of 1.00 for both gene signatures (Figure S4C).

#### Zscore vs. Original Model

The performance of gene signatures using Zscore was similar to GSVA, for which more datasets were selected for Anderson_OD_51, Berry_393, Berry_OD_86, Bloom_OD_144, and Zak_RISK_16. However, a distinct prediction pattern was visible when comparing these to their original methods (−0.0037 <= *ρ*_*w*_ <= 0.65, |∆*AUC*|_*w*_ > 0.05 for all five gene sets; Figure S4D). Similar to the results from ssGSEA and GSVA, Maertzdorf_15 (*ρ*_*w*_ = −0.85, |∆*AUC*|_*w*_ = 0.044) and Maertzdorf_4 (*ρ*_*w*_ = −0.74, |∆*AUC*|_*w*_ = 0.084) had negatively correlated prediction scores given using Zscore and their respective original models (Figure S4D).

#### Singscore unidirectional scoring vs. Original Model

Although more studies with high AUCs were selected when assessing Anderson_OD_51, Berry_393, Berry_OD_86, Bloom_OD_144, and Zak_RISK_16 with Singscore unidirectional scoring, results from these signatures had weak to moderate weighted Spearman’s rank correlation (0.16 <= *ρ*_*w*_ <= 0.55) and a large AUC difference (|∆*AUC*|_*w*_ >= 0.11 for all five gene sets; Figure S4E). Conversely, no datasets with high AUCs were selected for Anderson_42 (*ρ*_*w*_ = −0.23, |∆*AUC*|_*w*_ = 0.26) based on results given by Singscore unidirectional scoring. Sweeney_OD_3 (*ρ*_*w*_ = 0.85, p-value < 1e-04; |∆*AUC*|_*w*_ = 0.031, p-value = 0.75) was the only gene set for whom results from Singscore could act as a proxy to its original models, except for study GSE34608 (Figure S4E).

#### Gene signatures after splitting

Figure S5A-E shows the performance of 13 signatures as evaluated with their up/down regulated subset of genes using ssGSEA, GSVA, and Singscore bidirectional scoring. Notably, for the upregulated subset of Verhagen_10, none of the datasets had AUCs greater than 0.8, as given by ssGSEA and GSVA (Figure S5A-D). Study GSE101705 was the only selected study when evaluating the downregulated subset of Verhagen_10 using ssGSEA (*ρ*= 0.60, |∆*AUC*| = 0.20), GSVA (*ρ*= 0.57, |∆*AUC*| = 0.17), and Singscore bidirectional scoring (*ρ*= −0.63, |∆*AUC*| = 0.18; Figure S5B-E). However, it showed different prediction results than the original models. Additionally, GSE107994 was selected for the downregulated subset of Verhagen_10 given by GSVA (*ρ*= 0.44, |∆*AUC*| = 0.19; Figure S5D).

#### ssGSEA with signature split vs. Original Model

When assessing gene signatures based on their upregulated subset, all selected studies had positive Spearman’s rank correlations of the predicted scores given by ssGSEA and original models (Figure S5A). More studies with high AUCs were selected for 10 out of 12 gene signatures (excluding Leong_RISK_29 and Maertzdorf_15) when evaluated with their respective upregulated genes using ssGSEA. Results from subsets of Kaforou_27 and Kaforou_OD_53 showed similar diagnostic features when compared to their original models, with highly correlated prediction scores (*ρ*_*w*_ > 0.80 for both gene sets; p-value <= 1e-04) and small AUC differences (|∆*AUC*|_*w*_ < 0.030 for both gene sets; p-value > 0.080), and a study overlap coefficient of 1.00 for both gene sets (Figure S5A).

When gene signatures were evaluated using their downregulated subsets, only Maertzdorf_15 had an absolute Spearman’s rank correlation greater than 0.80 (*ρ*_*w*_: −0.89, p-value <= 1e-05) and a small AUC difference (|∆*AUC*|_*w*_ = 0.033, p-value = 0.010) for results given by ssGSEA and its original model, with a study overlap coefficient of 1.00 (Figure S5B). Only Berry_393 (*ρ*_*w*_ = −0.62), Leong_24 (*ρ*_*w*_ = −0.64), and Verhagen_10 (*ρ*_*w*_ = 0.60) had a more studies with high AUCs using ssGSEA (0.052 < |∆*AUC*|_*w*_ < 0.191; Figure S5B).

#### GSVA with signature split vs. Original Model

Figure S5C shows that the predicted scores given by GSVA and original models are positively correlated for most of the selected studies except GSE34608, for which the study was negatively correlated when evaluated by the upregulated subsets of Leong_24 (*ρ*= −0.11, |∆*AUC*| = 0.25). Moreover, Kaforou_27 had the highest weighted Spearman’s rank correlation (*ρ*_*w*_ = 0.90, p-value < 1e-05) and the smallest AUC difference (|∆*AUC*|_*w*_ = 0.020, p-value = 0.68) among 12 gene sets when its upregulated subset was assessed by GSVA, which presented an equivalent prediction pattern compared to its original model (Figure S5C). The upregulated subset of Maertzdorf_15 was the only gene set whose original model results outperformed GSVA, which had a greater number of studies with high AUCs from its original method (*ρ*_*w*_ = 0.68, |∆*AUC*|_*!*_ = 0.094; Figure S5C).

When gene signatures were evaluated with their downregulated subsets, the results from most of the gene sets given by GSVA underperformed the results from their original models. Specifically, nine of 13 gene sets had a greater number of studies with high AUCs based on the results from their original model (Figure S5D). Notably, no datasets with high AUCs were selected when the downregulated subset of Bloom_OD_144 and Kaforou_OD_44 were assessed by GSVA. GSE28623 was the only study with high AUCs for the down-regulated subset of Anderson_42 (Figure S5D). Maertzdorf_15’s GSVA results, similar to those for the ssGSEA method, had the highest absolute weighted Spearman’s rank correlation (*ρ*_*w*_ = −0.88, p-value <= 1e-05) and the smallest absolute AUC difference (|∆*AUC*|_*w*_ = 0.03, p-value = 0.20) when compared to its original model, with a study overlap coefficient of 1.00 (Figure S5D).

#### Singscore bidirectional scoring vs. Original Models

Most of the selected studies had positive correlations for the predicted scores given by Singscore bidirectional scoring and their original methods, except study GSE62525 from Bloom_OD_144 and GSE101705 from Verhagen_10 (Figure S5E). Both Kaforou_27 and Kaforou_OD_53 had a weighted Spearman’s rank correlation greater than 0.85 (p-value < 1e-05 for both gene sets), absolute AUC difference smaller than 0.020 (p-value > 0.05 for both gene sets), and study overlap coefficient of 1.00, which suggests that results for these gene signatures given by Singscore bidirectional scoring could act as a proxy from the results given by signatures’ original models (Figure S5E).

Signatures Anderson_OD_51, Kaforou_OD_44, Maertzdorf_15, and Sambarey_HIV_10 had a larger number of studies with high AUCs given by their original model, weighted Spearman’s rank correlations between 0.53 and 0.86, and absolute AUC differences ranging from 0.021 to 0.16 (Figure S5E).

## DISCUSSION

TB diagnostics is moving toward using blood-based biomarkers, but serious gaps remain in the analyses of these data. Here we evaluated the performance of 19 TB gene signatures in distinguishing active TB from other clinical conditions using the original published model and five different gene set scoring methods based on 24 transcriptomic studies. These datasets represent real-world heterogeneity concerning geographic regions, host and pathogen genetics, and clinical context (34, 35, 37). Our results suggested that an original gene signature model’s predictive ability can be improved or recaptured using gene set scoring methods.

The five gene set scoring methods used here belong to a class of methods that compute a gene set enrichment score for each sample using only the genes from a signature. However, some differences between methods are present. Gene set scoring methods, including ssGSEA, GSVA, Zscore, and Singscore are *single-sample* methods that rank genes in each sample individually by comparing the ranks of the signature genes with the ranks of non-signature genes in the sample. By design, using these single-sample methods, the score for an individual sample is not affected by other samples in the cohort. In contrast, in computing an individual sample score, PLAGE implements singular value decomposition (SVD) on the standardized gene expression profile of all subjects in the dataset (50, 51, 53) such that sample composition had a large effect on the PLAGE scores. In addition, the original models for the gene signatures Sweeney_OD_3, Kaforou (Kaforou_27, Kafotrou_OD_44, and Kaforou_OD_53), and Anderson (Anderson_42 and Anderson_OD_51) could also be characterized as single sample methods, which rely on the expression of upregulated and downregulated subset of genes within gene sets (Table 1). These single-sample methods were more likely to produce robust scores for individual subjects, especially in studies with small sample sizes or heterogeneous disease subtypes. Multi-sample methods were susceptible to sample composition (i.e., the number of samples for different subtypes), which could introduce biases if the size of samples for different disease subtypes changed (53). PLAGE and other multi-sample biomarker models (e.g., random forest, linear discriminant analysis, and K-nearest neighbors) compute a different score for an individual sample whenever a patient enters or leaves the test population. This is known as “test set bias” (62), as a gene signature’s performance may be irreproducible when a patient population composition or size changes. Therefore, one of the limitations of using these multi-sample methods for sample scoring is that bias will be induced if researchers perform integration analysis by merging datasets from different sources. In our study, the results given by PLAGE for most of the signatures outperformed the rest of gene set scoring methods (Table 3; Figure 2A). This is primarily because the gene signature selection process is usually heavily based on prior knowledge and the activity levels of the selected genes are biologically meaningful, which could be revealed using SVD-like methods, including principal component analysis (51).

The weighted mean AUCs given by single-sample scoring methods were sometimes lower than the AUC from the original model for some signatures (Table 3). For these cases, the biomarker splitting strategy improved diagnostic ability when using the single-sample scoring methods, ssGSEA, GSVA, and Singscore, consistent with existing publications in other fields (50, 63). Further, we noted that the improvement of weighted mean AUC values based on upregulated subsets was more dominant when compared to the results from the downregulated subset (Table 3). This is likely because upregulated genes are usually immune-related, such as FCGR1A/B, GBP5/6, C1QB, SEPTIN4, and ANDKRD22 (21), which generate a clear signal in active TB and are features of the immune response to the disease (64). The weighted AUCs from Zak_RISK_16 were consistently greater than 0.80 for five gene set scoring methods (Table 2), mainly due to the overexpression of all genes within Zak_RISK_16 relative to its discovery dataset, with large number of genes being highly differentially expressed from the recent Leicester clinical phenotype groups (64). Therefore, the choice of gene set scoring method depends on the cohort composition and the differential expressed genes within the gene signatures. Generally, it is ideal and efficient to implement gene set scoring methods to evaluate the diagnostic accuracy of TB signatures, which we have shown to achieve equivalent accuracy when compared to their original models.

The data preprocessing and training procedures are specialized and intractable for most TB gene signature discovery cases, which contributes to low generalizability in some biomarkers when evaluating their performance on independent datasets by implementing the original classification model. Both Berry_393 and Berry_OD_86 gene signatures used the K-nearest neighbor (KNN) algorithm, which demonstrated high classification ability in their discovery studies (Table 2) but had poor results across multiple studies (Figure 1). KNN clustering worked well when gene expression values were normalized to the median of each control group (65); however, performing such normalization for transcriptomic datasets originating from different clinical conditions or different platforms is unrealistic. KNN classification also assumes that similar inputs share similar labels; however, data points tend to be close together in high dimensional spaces (66). Since the Berry_393 and Berry_OD_86 consist of 393 and 86 transcripts respectively, KNN lost predictive power in transcriptomic studies containing few samples but large numbers of gene expression features.

On the other hand, the performance of Verhagen_10 was poor across independent studies based on the results given by its original diagnostic model but had AUC values equal to 1.00 in datasets GSE81746 and GSE41055 (Figure 1). GSE41055 is the discovery study for Verhagen_10, and GSE81746 is a small study that only contains two active TB diseases and four health control samples. In our comparison, 17 of 19 signatures could identify the active TB subtype with an AUC greater than 0.80 (Figure 1). The mediocre performance of Verhagen_10 across multiple studies is a sign of overfitting, a common problem using random forest, which relies on optimizing the tuning parameters (26). Since the original publication did not offer information about the hyper-parameters, we could not reconstruct the diagnostic model in the same way. Similar observations were made for Leong_RISK_29 (19), for which AUC results varied from 0.5 and 1.00 across independent datasets (Figure 1). As Leong_RISK_29 was designed to capture early host response to infection, it has not been generalized to samples with clinical manifestations of the disease (32). In these and many other ways, a gene signature’s diagnostic accuracy may be underestimated by evaluating its performance using its original model.

We also examined the gene signatures’ predictive accuracy within individual studies based on the results given by gene set scoring methods and the signatures’ original model. Results from PLAGE and ssGSEA for Kaforou_27 were highly correlated (*ρ*_*w*_ from PLAGE: = −0.92, *ρ*_*w*_ from ssGSEA: = 0.78), with a small AUC difference (|∆*AUC*|_*w*_ from PLAGE: = 0.030, |∆*AUC*|_*w*_ from ssGSEA: = 0.032), and it had a study overlap coefficient of 1.00 when compared to its original model (Figure S4A-B). This is primarily because of the presence of 21 overexpressed genes within the Kaforou_27 gene set, which dominates the performance of the gene signatures (Figure S5A). We identified a similar situation for Maertzdorf_15, which had 12 downregulated genes within its gene set. Maertzdorf_15 and its downregulated subsets had consistently high negative Spearman’s rank correlations (−0.89 <= *ρ*_*w*_ <= −0.78) for the prediction scores given by its original model and gene set scoring methods except for PLAGE (*ρ*_*w*_ = 0.94) and Singscore bidirectional scoring (*ρ*_*w*_ = 0.86; Figure S4 A-E; Figure S5B, D-E). The outperformance of Sweeney_OD_3 in ssGSEA and Singscore unidirectional scoring was largely due to the down-regulated gene *KLF2* within the Sweeney_OD_3 gene set. It has a relatively lower summary effect of the standardized mean difference between active tuberculosis and other clinical conditions from the biomarker’s original discovery datasets of Sweeney_OD_3; in contrast, the remaining two upregulated genes were highly differentially expressed in the corresponding datasets (26).

Our study has several limitations. We only compared the performance of active TB disease versus all other disease states regardless of how TB was diagnosed, and we did not perform subgroup analysis stratified by different clinical conditions, such as age, region, gender, or comorbidities. Additionally, some reconstructed signature models may not be the same as those used in the original publication, as some of these publications do not provide precise details to recreate the original training model. Instead, we followed the details outlined in the previously published comparison of Warsinske *et al*. (19). Due to different naming sequencing platforms for transcripts, some genes within signatures were missing across multiple studies, and we could not evaluate the biomarkers’ diagnostic ability in those cases accurately. Although we used KNN imputation to estimate the expression values for those missing genes in the validation study, the imputation process may have potentially led to bias. Our comparison would still be valid if we followed similar procedures to handle missing values across independent studies.

In conclusion, we showed that gene set scoring methods are effective for evaluating gene signature accuracy for comparing active TB disease versus other clinical conditions. These methods perform similarly when compared to the signatures’ original models. Overall, the performance of TB biomarkers across multiple studies was similar between the PLAGE and the original models. The methods ssGSEA, GSVA, and Singscore can also capture the diagnostic accuracy of gene signatures by taking the gene directional information within gene sets into account. Given the challenges associated with rebuilding or re-evaluating the signatures’ original biomarker model(s), gene set scoring methods could serve as a reliable alternative computational methodology to apply or perform comparisons of TB biomarkers.

---

**Supplementary Table 1.**
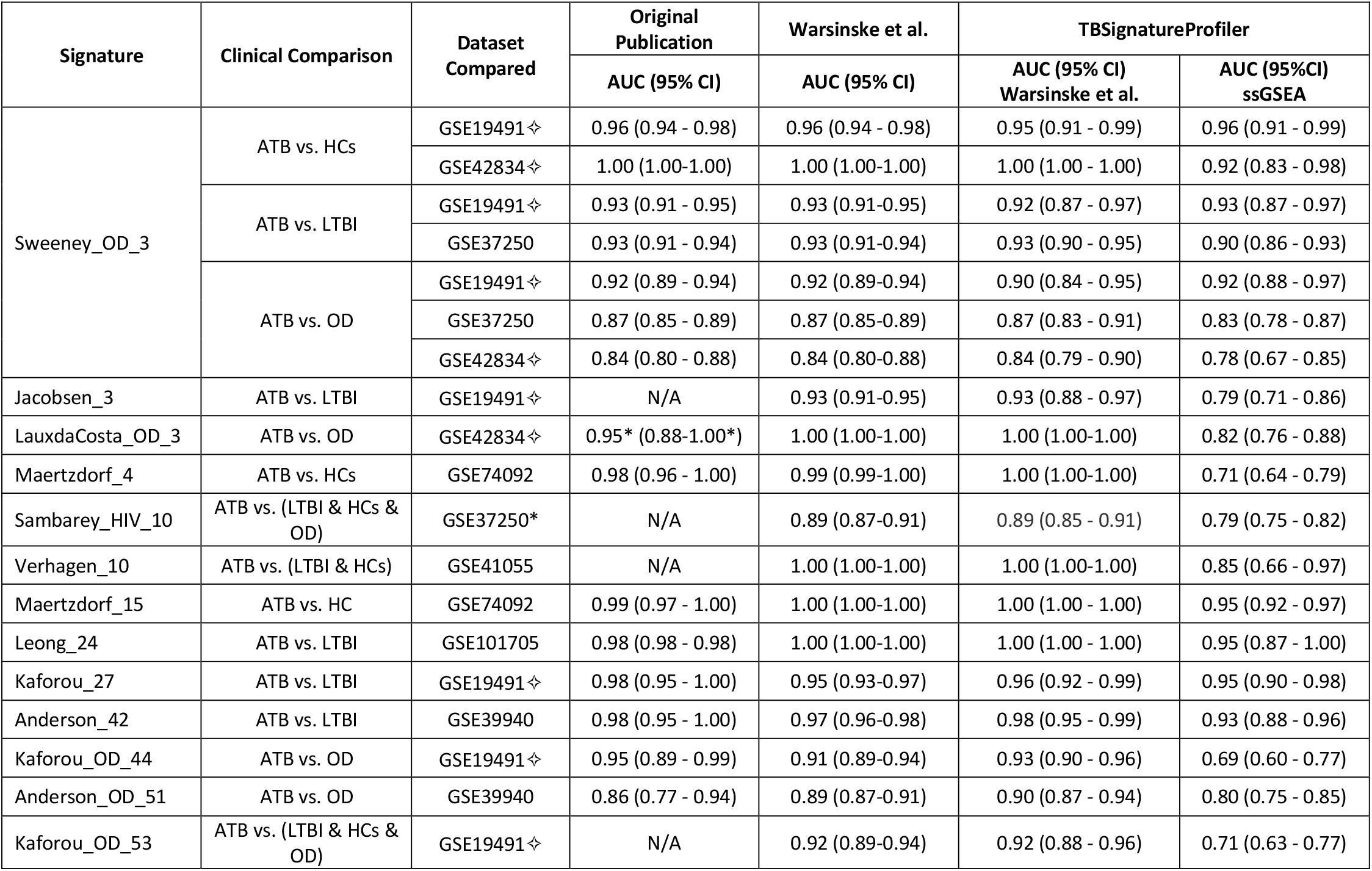
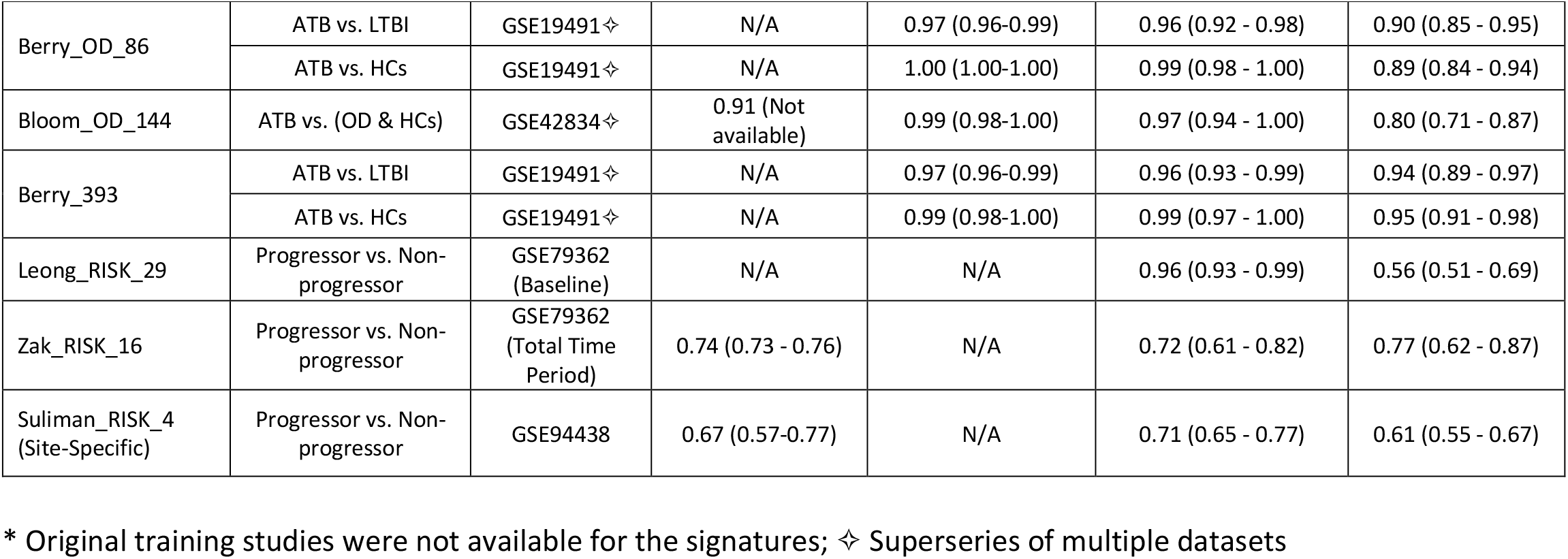
Summary of AUC and 95% CI results from gene signature’s discovery/training study. The original publication section presented the AUC and 95% CI for each signature from their publication. The Warsinske et al. subsection showed the results that have been re-evaluated using the original models functionality that had been implemented in the TBSignatureProfiler, and the ssGSEA section presented the performance of each gene signature using ssGSEA from TBSignatureProfiler (Courtesy to Warsinske et al. work; ATB: active tuberculosis).

| Signature | Clinical Comparison | Dataset Compared | Original Publication | Warsinske et al. | TBSignatureProfiler |  |
| --- | --- | --- | --- | --- | --- | --- |
|  |  |  | AUC (95% CI) | AUC (95% CI) | AUC (95% CI) Warsinske et al. | AUC (95%CI) ssGSEA |
| Sweeney_OD_3 | ATB vs. HCs | GSE19491↔ | 0.96 (0.94 - 0.98) | 0.96 (0.94 - 0.98) | 0.95 (0.91 - 0.99) | 0.96 (0.91 - 0.99) |
|  |  | GSE42834↔ | 1.00 (1.00-1.00) | 1.00 (1.00-1.00) | 1.00 (1.00 - 1.00) | 0.92 (0.83 - 0.98) |
|  | ATB vs. LTBI | GSE19491↔ | 0.93 (0.91 - 0.95) | 0.93 (0.91-0.95) | 0.92 (0.87 - 0.97) | 0.93 (0.87 - 0.97) |
|  |  | GSE37250 | 0.93 (0.91 - 0.94) | 0.93 (0.91-0.94) | 0.93 (0.90 - 0.95) | 0.90 (0.86 - 0.93) |
|  | ATB vs. OD | GSE19491↔ | 0.92 (0.89 - 0.94) | 0.92 (0.89-0.94) | 0.90 (0.84 - 0.95) | 0.92 (0.88 - 0.97) |
|  |  | GSE37250 | 0.87 (0.85 - 0.89) | 0.87 (0.85-0.89) | 0.87 (0.83 - 0.91) | 0.83 (0.78 - 0.87) |
|  |  | GSE42834↔ | 0.84 (0.80 - 0.88) | 0.84 (0.80-0.88) | 0.84 (0.79 - 0.90) | 0.78 (0.67 - 0.85) |
| Jacobsen_3 | ATB vs. LTBI | GSE19491↔ | N/A | 0.93 (0.91-0.95) | 0.93 (0.88 - 0.97) | 0.79 (0.71 - 0.86) |
| LauxdaCosta_OD_3 | ATB vs. OD | GSE42834↔ | 0.95* (0.88-1.00*) | 1.00 (1.00-1.00) | 1.00 (1.00-1.00) | 0.82 (0.76 - 0.88) |
| Maertzdorf_4 | ATB vs. HCs | GSE74092 | 0.98 (0.96 - 1.00) | 0.99 (0.99-1.00) | 1.00 (1.00-1.00) | 0.71 (0.64 - 0.79) |
| Sambarey_HIV_10 | ATB vs. (LTBI & HCs & OD) | GSE37250* | N/A | 0.89 (0.87-0.91) | 0.89 (0.85 - 0.91) | 0.79 (0.75 - 0.82) |
| Verhagen_10 | ATB vs. (LTBI & HCs) | GSE41055 | N/A | 1.00 (1.00-1.00) | 1.00 (1.00-1.00) | 0.85 (0.66 - 0.97) |
| Maertzdorf_15 | ATB vs. HC | GSE74092 | 0.99 (0.97 - 1.00) | 1.00 (1.00-1.00) | 1.00 (1.00 - 1.00) | 0.95 (0.92 - 0.97) |
| Leong_24 | ATB vs. LTBI | GSE101705 | 0.98 (0.98 - 0.98) | 1.00 (1.00-1.00) | 1.00 (1.00 - 1.00) | 0.95 (0.87 - 1.00) |
| Kaforou_27 | ATB vs. LTBI | GSE19491↔ | 0.98 (0.95 - 1.00) | 0.95 (0.93-0.97) | 0.96 (0.92 - 0.99) | 0.95 (0.90 - 0.98) |
| Anderson_42 | ATB vs. LTBI | GSE39940 | 0.98 (0.95 - 1.00) | 0.97 (0.96-0.98) | 0.98 (0.95 - 0.99) | 0.93 (0.88 - 0.96) |
| Kaforou_OD_44 | ATB vs. OD | GSE19491↔ | 0.95 (0.89 - 0.99) | 0.91 (0.89-0.94) | 0.93 (0.90 - 0.96) | 0.69 (0.60 - 0.77) |
| Anderson_OD_51 | ATB vs. OD | GSE39940 | 0.86 (0.77 - 0.94) | 0.89 (0.87-0.91) | 0.90 (0.87 - 0.94) | 0.80 (0.75 - 0.85) |
| Kaforou_OD_53 | ATB vs. (LTBI & HCs & OD) | GSE19491↔ | N/A | 0.92 (0.89-0.94) | 0.92 (0.88 - 0.96) | 0.71 (0.63 - 0.77) |

Supplementary Table 1 (Continued)
| Signature | Clinical Comparison | Dataset Compared | Original Publication | Warsinske et al. | TBSignatureProfiler |  |
| --- | --- | --- | --- | --- | --- | --- |
|  |  |  | AUC (95% CI) | AUC (95% CI) | AUC (95% CI) Warsinske et al. | AUC (95%CI) ssGSEA |
| Berry_OD_86 | ATB vs. LTBI | GSE19491✧ | N/A | 0.97 (0.96-0.99) | 0.96 (0.92 - 0.98) | 0.90 (0.85 - 0.95) |
|  | ATB vs. HCs | GSE19491✧ | N/A | 1.00 (1.00-1.00) | 0.99 (0.98 - 1.00) | 0.89 (0.84 - 0.94) |
| Bloom_OD_144 | ATB vs. (OD & HCs) | GSE42834✧ | 0.91 (Not available) | 0.99 (0.98-1.00) | 0.97 (0.94 - 1.00) | 0.80 (0.71 - 0.87) |
| Berry_393 | ATB vs. LTBI | GSE19491✧ | N/A | 0.97 (0.96-0.99) | 0.96 (0.93 - 0.99) | 0.94 (0.89 - 0.97) |
|  | ATB vs. HCs | GSE19491✧ | N/A | 0.99 (0.98-1.00) | 0.99 (0.97 - 1.00) | 0.95 (0.91 - 0.98) |
| Leong_RISK_29 | Progressor vs. Non-progressor | GSE79362 (Baseline) | N/A | N/A | 0.96 (0.93 - 0.99) | 0.56 (0.51 - 0.69) |
| Zak_RISK_16 | Progressor vs. Non-progressor | GSE79362 (Total Time Period) | 0.74 (0.73 - 0.76) | N/A | 0.72 (0.61 - 0.82) | 0.77 (0.62 - 0.87) |
| Suliman_RISK_4 (Site-Specific) | Progressor vs. Non-progressor | GSE94438 | 0.67 (0.57-0.77) | N/A | 0.71 (0.65 - 0.77) | 0.61 (0.55 - 0.67) |

\* Original training studies were not available for the signatures; ✧ Superseries of multiple datasets

**Supplementary Figure 1.**
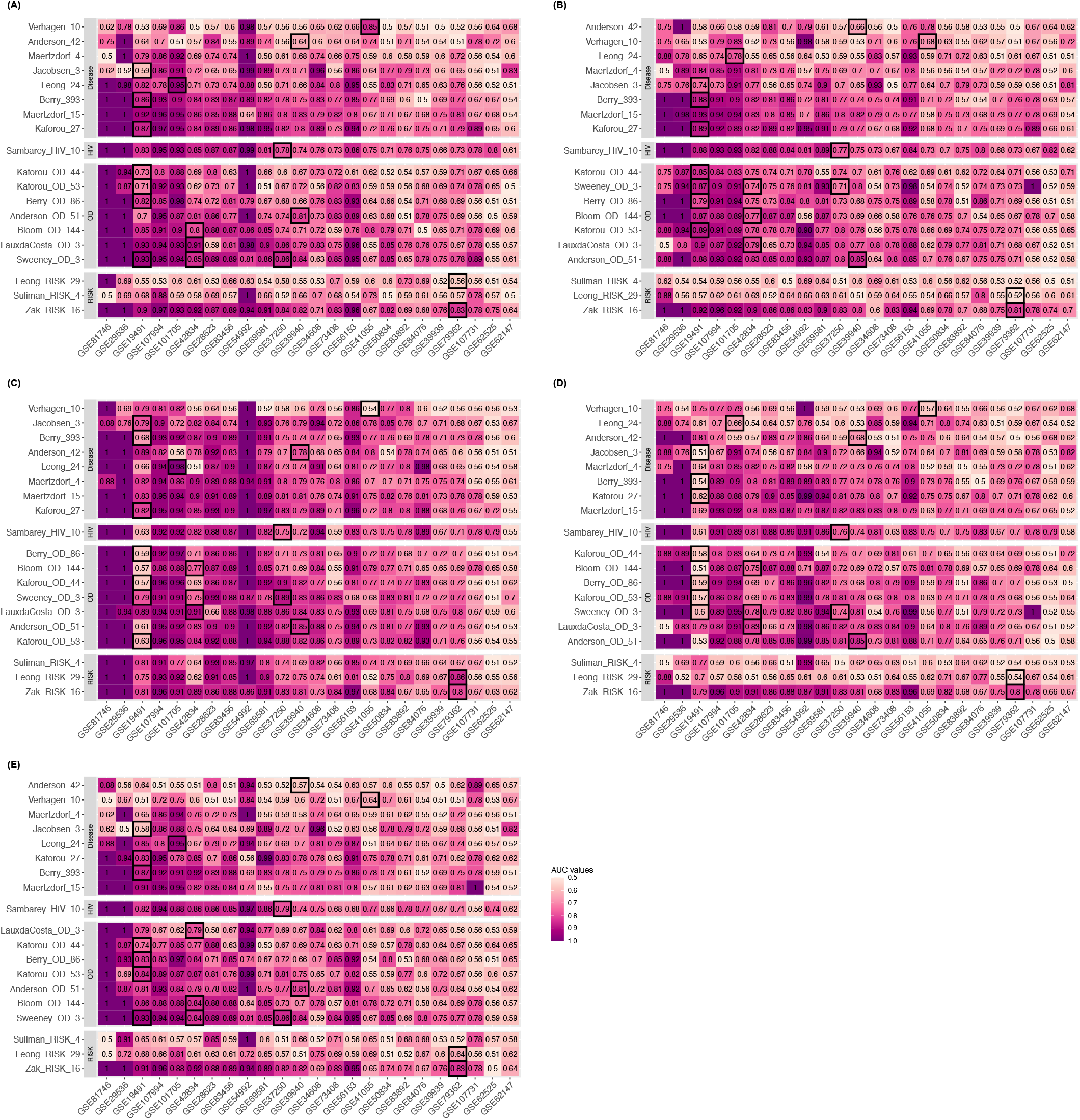
The distribution of AUC values for 19 TB gene signatures across 24 studies using ssGSEA **(A)**, GSVA **(B)**, PLAGE **(C)**, Zscore **(D)**, and Singscore unidirectional version **(E)**. Datasets had the same order as those from Figure 1.

**Supplementary Figure 2.**
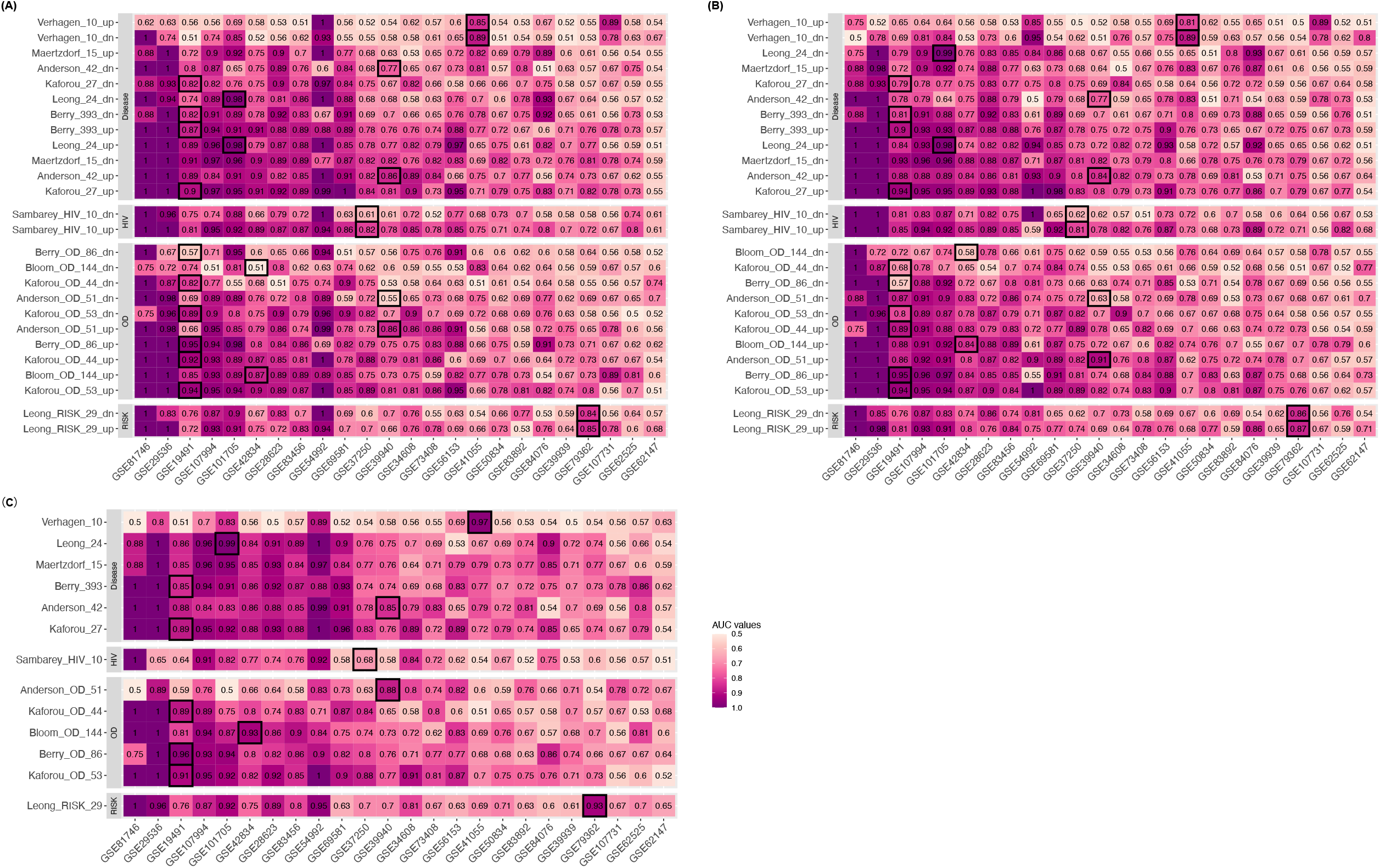
The distribution of AUC values for **upregulated and downregulated subsets** of 13 TB gene signatures across 24 studies using ssGSEA **(A)**, GSVA **(B)**, and Singscore bidirectional version **(C)**. Datasets had the same order as those from Figure 1.

**Supplementary Figure 3.**
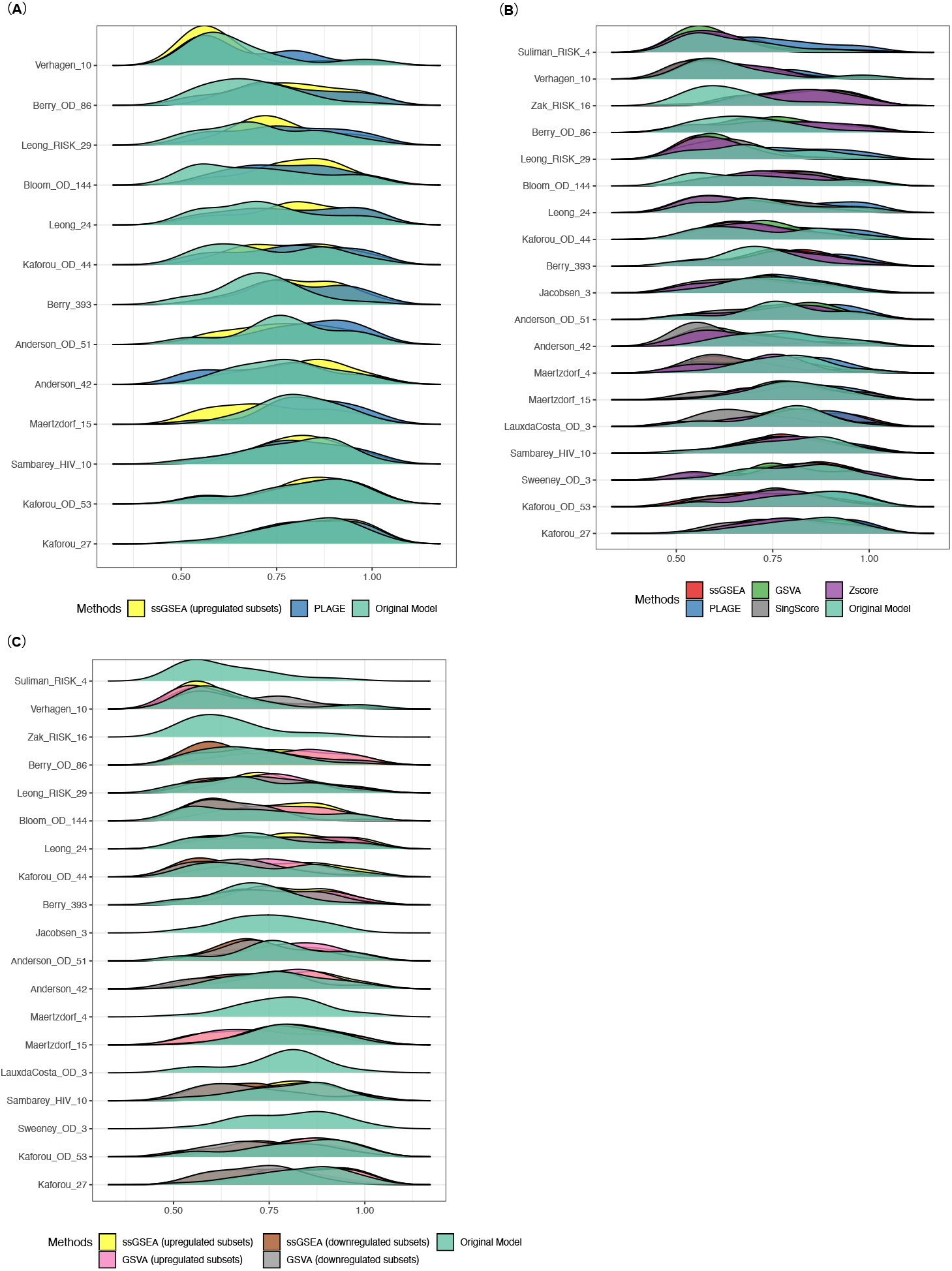
The distribution of AUC values for TB gene signatures across 24 studies using the original models and gene set scoring methods in the form of ridge plot. Gene signatures were ordered based on the median AUCs given by the original models. **(A)** Comparison of AUC distributions given by ssGSEA (evaluated on the upregulated subsets of the gene signatures), PLAGE, and the original models. **(B)** Comparison of AUC distributions given by five gene set scoring methods (ssGSEA, PLAGE, GSVA, Singscore, Zscore) and the original models. **(C)** Comparison of AUC distributions given by ssGSEA (evaluated on the upregulated and downregulated subsets of gene signatures), GSVA (evaluated on the upregulated and downregulated subsets of gene signatures), and the original models.

**Supplementary Figure 4.**
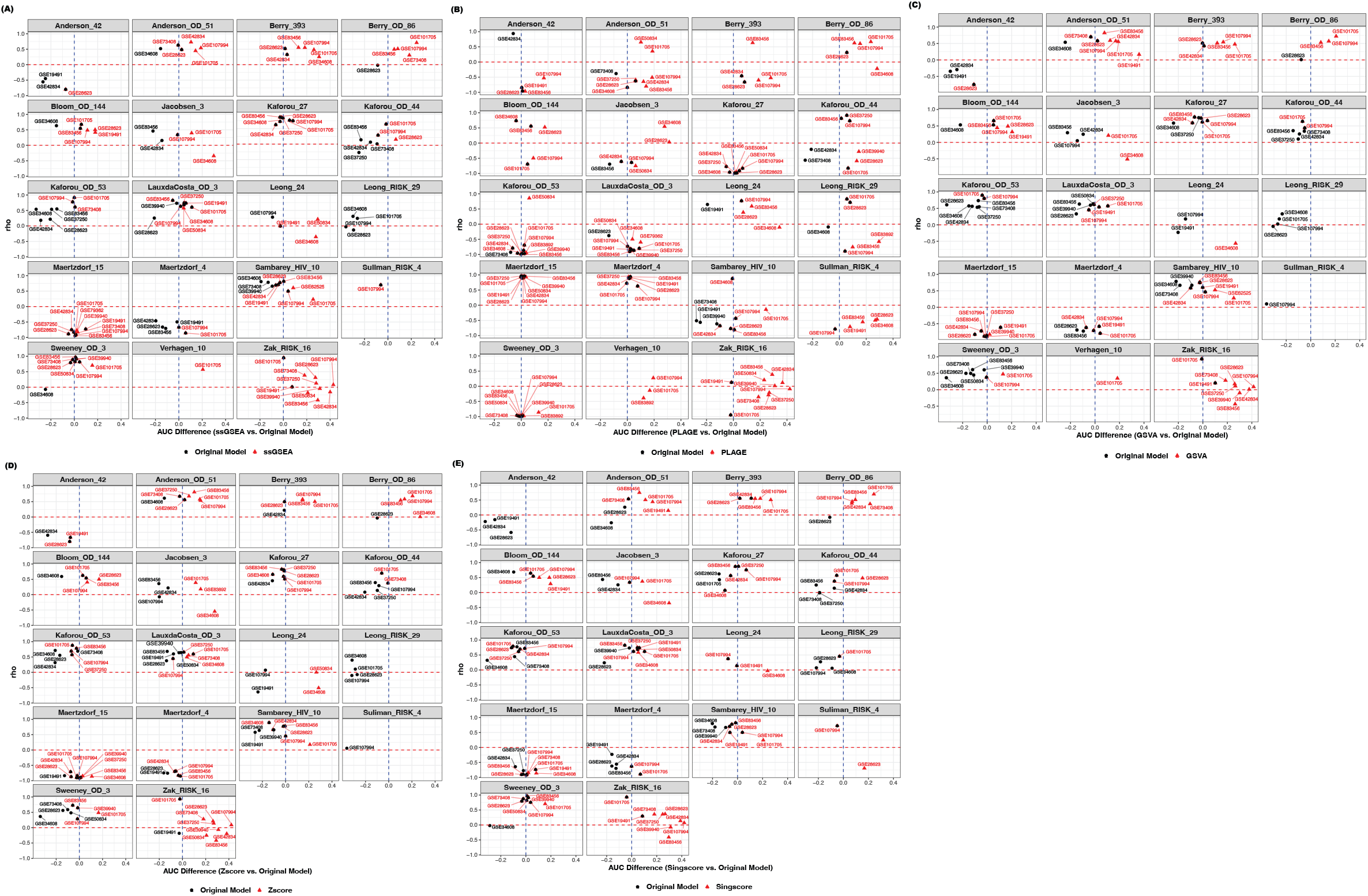
Spearman’s rank correlation versus AUC difference for studies with sample size larger than 40 and computed AUC values greater than 0.8 based on the results given by original models and ssGSEA **(A)**, original models and PLAGE **(B)**, original models and GSVA **(C)**, original models and Zscore **(D)**, original models and Singscore (unidirectional version) **(E)**. Overlapping points indicate the same dataset(s).

**Supplementary Figure 5.**
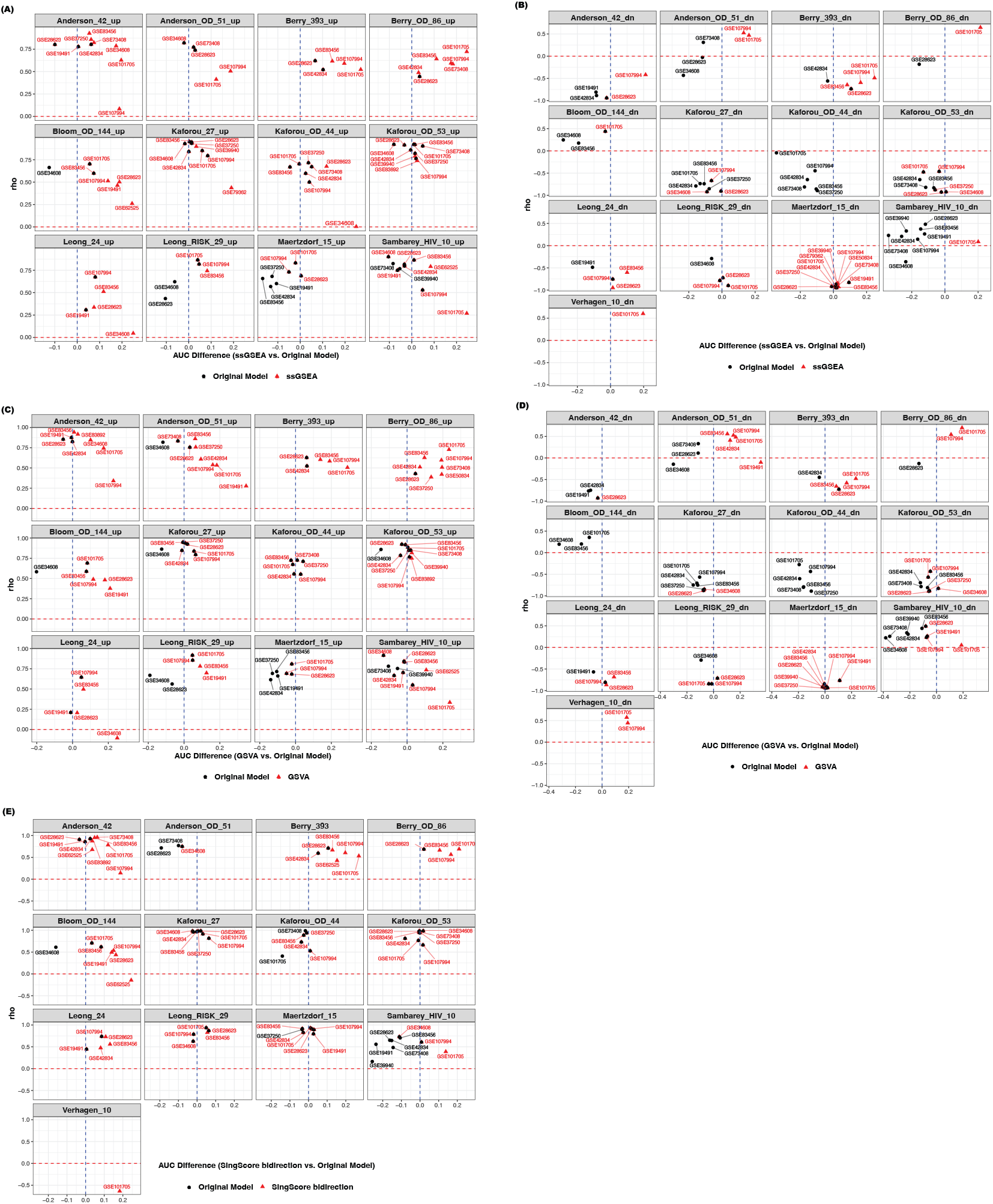
Spearman’s rank correlation versus AUC difference for studies with sample size larger than 40 and computed AUC greater than 0.8 based on results given by original models and ssGSEA (evaluated with upregulated subsets) **(A)**, original models and ssGSEA (evaluated with downregulated subsets) **(B)**, original models and GSVA (evaluated with upregulated subsets) **(C)**, original models and GSVA (evaluated with downregulated subsets) **(D)**, original models and Singscore (bidirectional version) **(E)**. Overlapping points indicate the same dataset(s).

